# Mechanism and kinetics of copper complexes binding to the influenza A M2 channel

**DOI:** 10.1101/2020.08.24.265165

**Authors:** K. McGuire, P. Smit, D. H. Ess, J. T. Hill, R. G. Harrison, D. D. Busath

## Abstract

Copper(II) is known to bind in the influenza virus His37 cluster in the homotetrameric M2 proton channel and block the proton current needed for uncoating. Copper complexes based on iminodiacetate also block the M2 proton channel and show reduced cytotoxicity and zebrafish-embryo toxicity. In voltage-clamp oocyte studies using the ubiquitous amantadine-insensitive M2 S31N variant, the current block showed fast and slow phases in contrast to the single phase found for amantadine block of WT M2. Here we evaluate the mechanism of block by copper adamantyl iminodiacitate (Cu(AMT-IDA)) and copper cyclooctyl iminodiacitate (Cu(CO-IDA)) complexes and address whether the complexes can covalently bind to one or more of the His37 imidazoles. The current traces were fitted to parametrized master equations. The energetics of binding and the rate constants suggest that the first step is copper-complex binding within the channel and the slow step in the current block is the covalent bond formation between copper complex and histidine. Isothermal titration calorimetry (ITC) indicates that a single imidazole binds strongly to the copper complexes. Structural optimization using density functional theory (DFT) reveals that the complexes fit inside the channel and project the Cu(II) towards the His37 cluster allowing one imidazole to form a covalent bond with the Cu(II). Electrophysiology and DFT studies also show that the complexes block the G34E amantadine-resistant mutant in spite of some crowding in the binding site by the glutamates.

## Introduction

Influenza A virus pandemics of zoonotic origin occur every few decades and cause respiratory disease globally. The influenza A M2 channel is responsible for the acidification of the influenza virus interior, which leads to the separation of ribonuclear proteins from the virus, at which point replication begins. Protons are transported to the viral interior through the protonation of imidazole nitrogens in the His37 cluster (Figure 1). Amantadine and rimantadine block the M2 WT variant but they do not block the ubiquitous M2 S31N variant, nor other naturally occurring variants including V27A, A30T, and G34E (1), and are no longer FDA-approved as influenza A anti-viral drugs. Numerous studies have been carried out in search of organic compounds to block these mutants (2–9). None has been shown to be effective against all naturally occurring influenza A M2s (10), although a few compounds have recently been identified that can block some of the most common single and double mutants (11).

**Figure 1:**
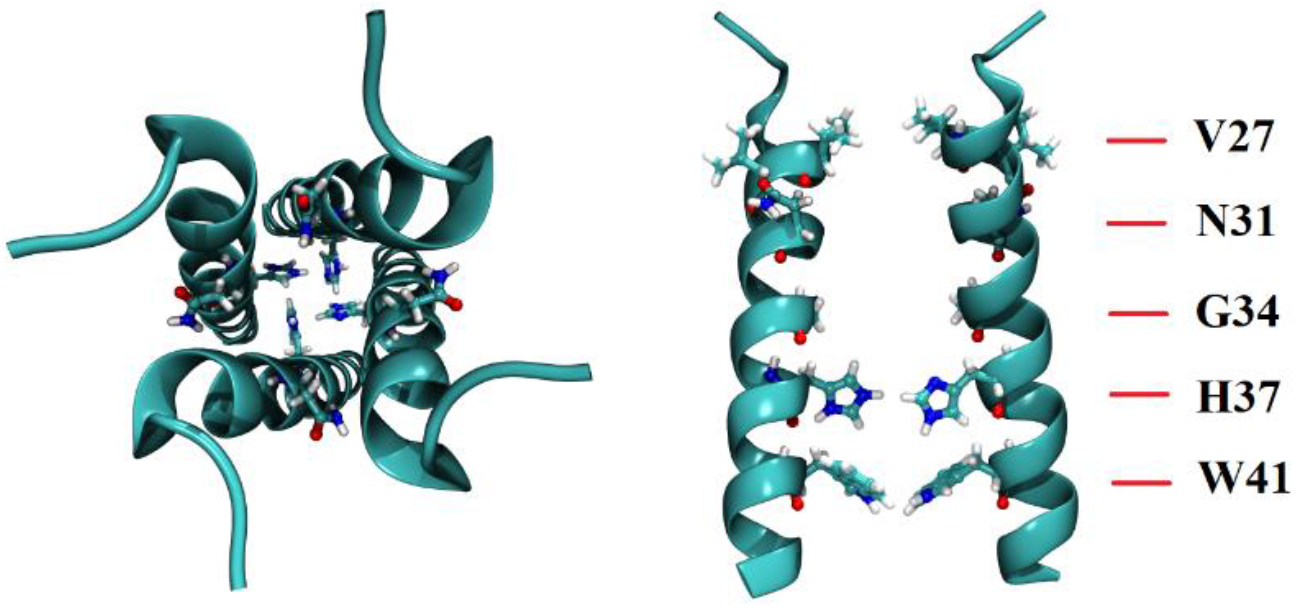
M2 S31N ssNMR Structure (2KQT). The model shows the TM domain (residues 22-46). The left image is a top-down view (N-terminus to C-terminus). The right image is a side-view with two of the peptide backbones and the side chains hidden.

The His37 cluster is an excellent target for an antiviral because it is almost perfectly conserved in nature (12), probably because of its key role in selecting (13) protons for transport into virions after endocytosis. The His37 cluster was shown to be a viable binding site for copper complexes (12) based on the knowledge that Cu^2+^ binds to imidazole (14) and blocks the M2 channel (15) from a position between the His37 and Trp41 clusters (16). Previous work using the electrophysiology two-electrode voltage method (TEVC) showed that Cu(CO-IDA) and Cu(AMT-IDA) block the M2 S31N variant at low (therapeutic) concentrations similar to AMT block in M2 WT. A two-phase block was observed, an initial fast phase followed by a slow phase (12). By site-directed mutagenesis of the H37 to A37, the complexes were shown to interact with the His37 cluster. The mutagenesis resulted in elimination of the slow phase, leaving the initial fast phase of block followed by complete wash-out in complex-free perfusate (12).

*In vitro,* Cu(CO-IDA) and had low cytotoxicity and was potent at sub-micromolar concentrations against A/Calif/07/2009 H1N1, which contains M2 S31N (17). In an evaluation of M2 resistance development against these copper complexes, 10 passages were insufficient for resistance development in contrast to resistance to amantadine, which develops after only a few passages (17). However, there is another naturally occurring mutation, G34E, that could interfere with binding of the copper complexes because of the proximity to the binding site and added bulk of the glutamate side chains. Although this mutation did not appear spontaneously during virus passaging, we thought it prudent to test the copper complexes in the M2 S31N-G34E variant using TEVC, and to explore its possible copper complex binding configurations with DFT.

This research explores the mechanism by which Cu(CO-IDA) and Cu(AMT-IDA) block the M2 channel. We explore the binding kinetics and configurations of the copper complexes with M2 S31N. A global fit of electrophysiology data allows the determination of rate constants, providing insight into the association, covalent binding, and dissociation processes. Isothermal titration calorimetry (ITC) was used to evaluate the enthalpy of bonding and dissociation constant for the binding of imidazole to the copper complexes. Quantum mechanical simulations help elucidate the configuration of the copper complexes bound to histidines in the channel.

## Methods and Materials

### A/Udorn/72 H3N2 M2 S31N-G34E mRNA Synthesis

The preparation of mRNA for oocytes with the influenza A/Udorn/72 H3N2 M2 S31N protein were reported previously (12). Here we modified that plasmid to add a G34E mutation. A DNA fragment containing a G to E mutation to create the G34E allele was obtained from Twist Biosciences (San Francisco, CA). The fragment was digested using BamHI and HindIII restriction nucleases and cloned into the A/Udorn/72 H3N2 M2 S31N plasmid, digested with the same restriction nuclease, to create the A/Udorn/72 H3N2 M2 S31N-G34E plasmid. The plasmid was transformed into chemically competent E. coli by standard methods. The plasmid was harvested using the Zymo Miniprep Kit (Zymo Research, Irivine, CA). To confirm that no mutations were introduced, the M2 DNA segment was PCR amplified and Sanger sequenced (Figure S1). Following confirmation, the PCR product was transcribed using the mMESSAGE mMACHINE T7 ULTRA Transcription Kit (Thermo Fisher Inc., Waltham, MA) to prepare mRNA for oocyte injections.

### Electrophysiology

Here, we extend the experiments reported previously (12) by adding higher concentrations of metal complexes, normalizing each trace after a limited leak subtraction, and averaging together such traces from three separate experiments. For this, Cu(Amt-IDA)·5H_2_O (418.93 g/mol) and Cu(CO-IDA)·3H_2_O (358.88 g/mol) were synthesized according to previously published procedures (12). Oocytes from *Xenopus* laevis (Ecocyte, Austin, TX) were maintained in ND-96^++^ solution (96 mM NaCl, 2 mM KCl, 1.8 mM CaCl_2_, 1 mM MgCl_2_, 2.5 mM sodium pyruvate, 5 mM HEPES-NaOH, pH 7.4) at 17° C until injection of ~40 ng of A/Udorn/72 H3N2 M2 S31N or A/Udorn/72 H3N2 M2 S31N-G34E mRNA using a Nanoject II (Drummond Scientific, Broomall, PA). After injection, the oocytes were maintained at 4° C in ND96^++^ pH 7.4 until electrophysiological recording. Seventy-two hours after mRNA injection, whole-cell currents were recorded with a two-electrode voltage-clamp (TEVC) apparatus at a membrane potential Vm = −20 mV, room temperature, in Barth’s solution (0.3 mM NaNO_3_, 0.71 mM CaCl_2_, 0.82 mM MgSO_4_, 1.0 mM KCl, 2.4 mM NaHCO_3_, 88 mM NaCl, 15.0 mM HEPES, pH 7.5). Inward current was induced by perfusion with Barth’s pH 5.3 (15.0 mM MES instead of 15.0 mM HEPES). MES was chosen as a Good buffer for non-interaction with Cu(II) (18). The oocytes were then perfused by Barth’s pH 5.3 with Cu^2+^(aq), Cu(CO-IDA), or Cu(AMT-IDA) at concentrations 0.1 mM, 0.5 mM, or 1 mM. A wash-out was done using Barth’s pH 5.3 without complex. Non-injected oocytes were also tested with the same acid perfusion protocols and concentrations of complex to assess the possible native acid-activated channel and Cu^2+^(aq), Cu(CO-IDA), or Cu(AMT-IDA)-induced leak current in the oocytes. Leak-current subtracted current traces were obtained from each of three oocytes for each of the concentrations, normalized, and averaged.

### Global Nonlinear Least Squares Curve Fit

The two-phase block in the current traces (i.e. fast and slow phase) produced by perfusion with Cu^2+^(aq), Cu(CO-IDA) or Cu(AMT-IDA) were analyzed using the serial two-site binding model:

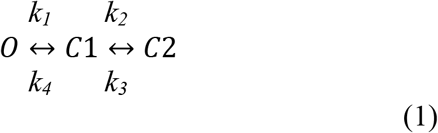

The open state, O, is the unblocked M2 channel where proton conductance is unimpeded prior to perfusion with blocking agents. C_1_ represents the initial bound state. Because the fast block never reaches completeness before the slow block begins, the permeability of C1 cannot be directly ascertained. We assume that the complex only partially blocks proton current in this state and we represent the fraction of current for this state with the fitting parameter *f*. Occupancy of the complex in the second binding site is represented by state C2, which is assumed to be a fully blocked M2 state because at high concentrations and long exposure full block is achieved. To obtain the rate constants, *k_i_*, they and *f* were used as parameters in Eq. 2 to fit the compound wash-in and wash-out current traces, similar to the method used to fit adamantanamine block in M2 WT and S31N currents (19).

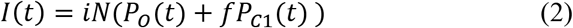

The single channel current at the applied membrane potential is *i* and the number of channels in the cell membrane is *N*. During drug wash-in, *I(t)*, the M2 S31N oocyte current, starts at a maximal level because the probability of being in the open state, *P_O_(0)* is 1 and declines with time as the sum of two exponentials towards a steady state level of block as the drug binding in the two states reaches equilibrium. The probabilities of the other two occupancy states are also functions of time, starting at no occupancy (*P_C1_(0)* = *P_C2_(0)* = 0) and approaching their equilibrium levels with a difference that also falls as the sum of two exponentials. During wash-out, the process is reversed and all three state probabilities relax back to their pre-drug occupancies as the sum of two exponentials. The underlying differential equations governing the relaxation of the probabilities following a change in boundary condition (bath drug concentration) and their solutions are given for both wash-in and wash-out in Supplemental. The rate constants and coefficients of the exponentials differ with drug concentration, and for the wash-out the coefficients also depend on the state probabilities achieved during the finite wash-in. Nonlinear least-squares curve fitting was carried out in MATLAB R2018a (The MathWorks Inc., Natick, MA) using the Levenberg-Marquardt algorithm. Uncertainties in the parameters were derived from the error matrix, i.e. the square roots of the diagonal elements (20).

To understand the strength of the interaction between M2 and the copper complexes, the effective equilibrium dissociation constant (dissociation from the second binding site to the open state, O) was calculated as the product of the outer and inner dissociation constants, *K_d1_* and *K_d2_*, using Eq. 3:

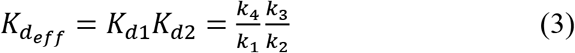

### Isothermal Titration Calorimetry

For ITC, Tris-buffered solutions of imidazole were prepared by adding imidazole to 8.0 mL of Milli-Q water, adding 1.0 mL of 20 mM Tris, adjusting the pH by adding either 10.0 or 1.0 M NaOH or HCl, and adding water to reach 10.0 mL of solution. Solutions of complexes were made by adding 41.5 mg Cu(Amt-IDA) or 27.3 mg Cu(CO-IDA) to 8 mL of Milli-Q water, adding 1.0 mL of 20 mM Tris, and sonicating and/or heating, until the complex dissolved (20-60 minutes). The pH was then adjusted to the target pH with 10.0 M or 1.0 M HCl or NaOH, and water added to reach 10 mL. Concentrations of copper complexes were verified by measuring UV absorbances of the complexes and using the Beer-Lambert law, and by ICP analysis. The molar extinction coefficient for Cu(Amt-IDA) is 5300 M^−1^cm^−1^ and for Cu(CO-IDA) is 3700 M^−1^cm^−1^ at 252 nm (12).

Isothermal titration calorimetry (ITC) experiments were performed using a Nano-ITC low-volume calorimeter (TA Instruments, Layton, UT) equipped with gold reference and sample cells of 170 μL, having a minimal detectable heat of 0.05 μJ and a baseline stability of .02 μW/hr. All titration runs were carried out with a 50 μL injection syringe (minimum injection volume 0.06 μL) at 25 °C and a stirring rate of 350 rpm. Both the syringe and well solutions were adjusted to the desired pH of 6, 7 or 8 using HCl or NaOH. One or two μL volumes of imidazole solution were injected into ~1 μM copper complex solutions (Cu(Amt-IDA) and Cu(CO-IDA)). Areas under the heat-of-injection curve for each experiment were corrected for heat of dilution, fitted using the *independent fit* in the NanoAnalyze software (TA Instruments, New Castle, DE), and fit parameters were averaged. The number of experiments for each complex ranged from 2 to 6. *K_d_* values depend primarily on the slope of the titration curve, *ΔH* on the intercept, and n, the number of imidazole binding sites on the complexes, modulated both. The value of n showed no pattern with complex or pH, and were therefore averaged over all fits, giving, *n* = 1.01 ± 0.37. On the grounds that variations in n probably reflected experimental fluctuations in complex integrity or solubilization, the modulating effects of *n* on *ΔH* and *K_d_* were incorporated in their averages. The first heat-of-injection peak in an experiment was routinely discarded due to dilution of the injectant by diffusion. Additionally, some experiments were rejected because of drifting baseline temperature or evidence of solution contamination due to incomplete cleaning of the calorimeter cell.

### Quantum Chemical Modeling of Cu(Amt-IDA) and Cu(CO-IDA)

As a quantum-chemical model for M2 channel binding, DFT calculations were used to examine the coordination enthalpy of neutral imidazole to Cu(CO-IDA). These calculations were performed using Terachem (v1.93P, PetaChem, LLC, Los Altos Hills, CA) using the ωB97X-D3 functional (21). Calculations were carried out with the default Terachem COSMO continuum solvent model (22). All structures were optimized to stationary points and confirmed as minima by vibrational frequency analysis using ωB97X-D3/6-31G** with the LANL2DZ used for copper. A subsequent SCF energy evaluation was performed with ωB97X-D3/6-311+G**, using LANL2DZ for copper, so that final enthalpy values, which contain the Δ*G* of solvation estimate, are ωB97X-D3/6-31G**[LANL2DZ Cu]//ωB97X-D3/6-311+G**[LANL2DZ Cu].

### Quantum Chemical Modeling – Copper Complexes Binding His37 in S31N and S31N-G34E M2

Previously, molecular dynamics simulations illustrated the accessibility of the channel lumen to these metal complexes (17). Here we explore whether a binding configuration with covalent bonding to a His37 side chain is feasible. Geometry optimization of the copper complexes binding to one histidine in the His37 cluster of the M2 S31N or S31N-G34E structure were done using Gaussian 16 (Gaussian, Inc., Wallingford, CT). A homology model for the M2 S31N and S31N-G34E was created from the 2KQT NMR structure with the VMD Mutator plugin. The M2-copper complex system was optimized in gas phase. ONIOM (23) was set up with the high-level QM region including all four histidines in the His37 cluster, Cu(AMT-IDA) or Cu(CO-IDA), and all four glycine or glutamate sidechains at positions 34. The low-level region included all other regions of the M2 channel. The ωB97X-D DFT method and 6-31G** basis set was used to optimize the QM region, except for the copper atom where the LANL2DZ basis was applied. The PM6 semi-empirical method was used to optimize the low-level regions.

## Results and Discussion

### Binding Kinetics

Binding kinetics were determined from electrophysiology block data for M2 S31N perfused with various concentrations of the three blockers (Figure 2). The normalized current traces are point-wise averages from three different *Xenopus* laevis cells with leak current subtracted from each trace. The simultaneous optimization in the global fit of the parameters from the block and wash-out electrophysiology data at varying concentrations constrained the parameters sufficiently to extract the rate constants (*k_1_, k_2_, k*_3_, and *k_4_*) and the fractional block, *f*, of the first binding site. For all three blockers, *f* was optimized at ≤ 0.02, i.e. complete block in the C1 state‥

**Figure 2:**
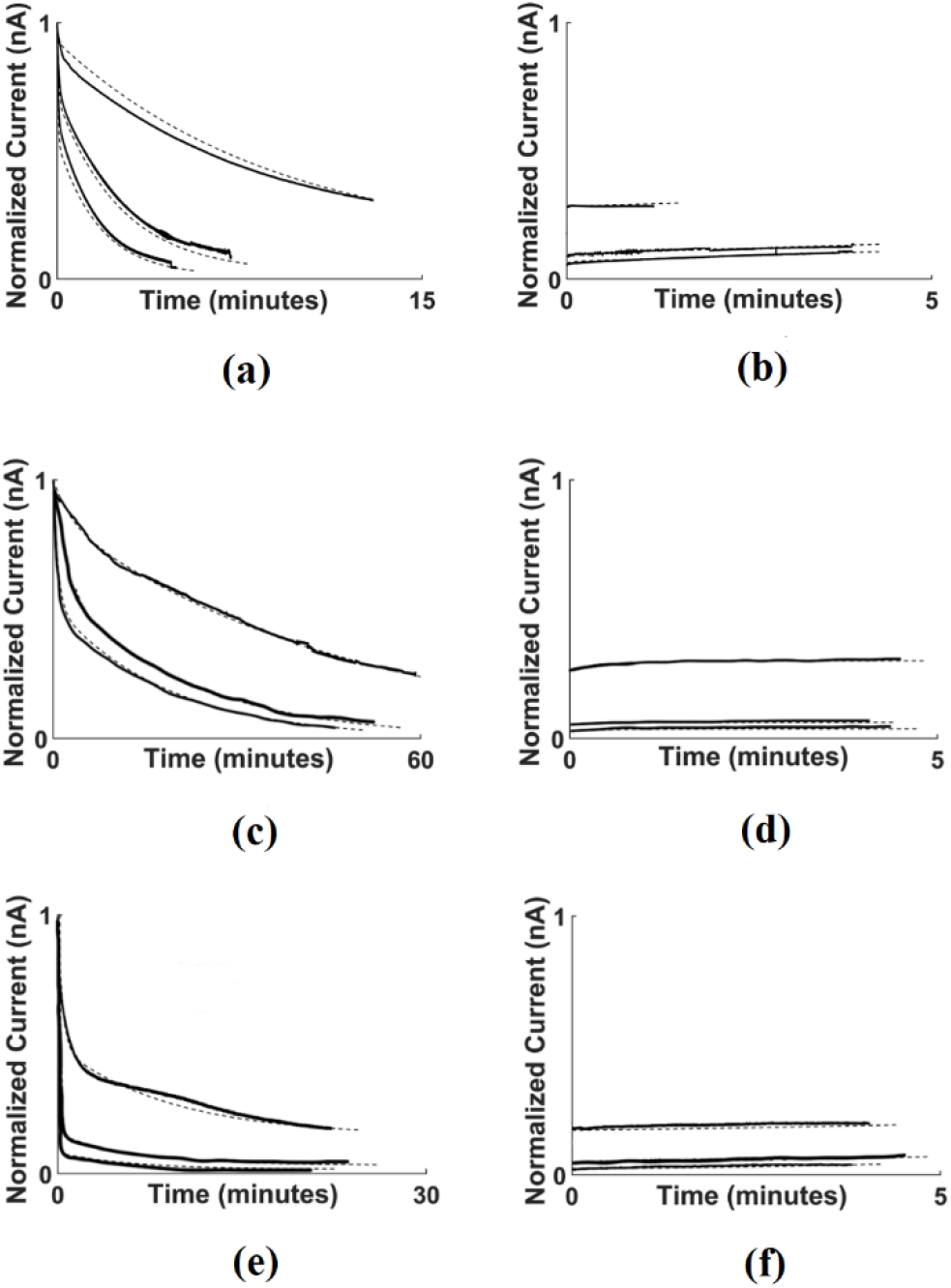
Cu2+(aq) (a,b), Cu(AMT-IDA) (c,d) and Cu(CO-IDA) (e,f), in S31N M2 tested at 0.10 0.50 and 1.0 mM. Perfusion of drug solution (wash-in) (a, c, and e) Drug-free perfusion (wash-out) (b, d, and f).

At 1.0 mM Cu^2+^(aq) the M2 S31N current is almost completely blocked in 5 minutes (Figure 2a). No significant wash-out was observed for Cu^2+^(aq) after 4 minutes (Figure 2b). The rate constants obtained in the slightly over-constrained global fit from the blocking and wash-out traces are 754 M^−1^ s^−1^ (*k_1_*) and 0.0824 s^−1^ (*k_2_*) for the binding to the first and second sites respectively, and 3.6×10^−4^ s^−1^ (*k_3_*) and 0.97 s^−1^ (*k_4_*) for the dissociation from the second and first binding sites respectively (Table 1). The corresponding effective equilibrium constant for the dissociation reaction of Cu^2+^(aq) from M2 S31N, *K_d,eff_*, is 56 nM (Table 2). This very low dissociation constant is primarily due to the strong binding at the second site. We propose the strong binding is due to the formation of a covalent bond between Cu(II) and a histidine residue in the channel.

**Table 1.** Association and dissociation rate constants for the first binding site (k_1_ and k_4_) and the second binding site (k_2_ and k_3_) from the global nonlinear least squares fit for Cu^2+^(aq), Cu(CO-IDA), and Cu(AMT-IDA) in M2 S31N.

| Global Fit Results |  |  |  |  |  |
| --- | --- | --- | --- | --- | --- |
| Compound | $k_1$ (M<sup>-1</sup>s<sup>-1</sup>) | $k_2$ (s<sup>-1</sup>) | $k_3$ (x10<sup>-5</sup> s<sup>-1</sup>) | $k_4$ (s<sup>-1</sup>) | Weighted Reduced $\chi^2$ |
| Cu<sup>2+</sup>(aq) | 754 ± 5 | 0.082 ± 0.012 | 0.36 ± 0.08 | 0.97 ± 0.09 | 0.00174 |
| Cu(CO-IDA) | 2876 ± 3 | 0.029 ± 0.004 | 2.9 ± 0.4 | 0.29 ± 0.01 | 0.00192 |
| Cu(AMT-IDA) | 594 ± 4 | 0.032 ± 0.003 | 1.5 ± 0.1 | 0.17 ± 0.01 | 0.00104 |

**Table 2.** Dissociation constants for the first and second binding steps and for the holistic binding process, i.e. from bulk to covalently bound state. Errors are propagated from those in Table 1 assuming no correlations.

| **Compound** | $K_{d1} (M) = k_4/k_1$ | $K_{d2} = k_3/k_2$ | $K_{d,eff}(M) = K_{d1} \times K_{d2}$ |
| --- | --- | --- | --- |
| Cu<sup>2+</sup>(aq) | $1.29 \times 10^{-3} \pm 1.06 \times 10^{-4}$ | $4.37 \times 10^{-5} \pm 1.13 \times 10^{-5}$ | $5.62 \times 10^{-8} \pm 1.53 \times 10^{-8}$ |
| Cu(CO-IDA) | $1.01 \times 10^{-4} \pm 3.83 \times 10^{-6}$ | $1.00 \times 10^{-3} \pm 1.98 \times 10^{-4}$ | $1.01 \times 10^{-7} \pm 2.03 \times 10^{-8}$ |
| Cu(AMT-IDA) | $2.86 \times 10^{-4} \pm 2.36 \times 10^{-5}$ | $4.69 \times 10^{-4} \pm 5.67 \times 10^{-5}$ | $1.34 \times 10^{-7} \pm 1.96 \times 10^{-8}$ |

Cu(AMT-IDA) blocks M2 S31N (Figure 2c) and produces blocking kinetics similar to Cu^2+^(aq). Almost complete block was achieved for 0.50 and 1.0 mM solutions by 60 minutes (Figure 4a) and no significant wash-out was observed by 5 minutes (Figure 2d). However, there are some significant differences in the fitted parameters with *k_1_* and *k_2_* lower, while *k_3_* is higher (Table 2), corresponding to a larger, yet still high-affinity value for *K_d,eff_*, 134 nM (Table 3). The increase in *k_3_* is consistent with binding being destabilized by configurational limitations imposed by the AMT-IDA ligand. From the low effective dissociation constant, and more particularly from the low value for the rate of dissociation from C2 (*k_3_* = 1.5×10^−5^ s^−1^), we propose that the copper complex also forms a covalent bond with the channel.

**Table 3:** The heats of complexation (Δ*H*) and binding constants (K_d_) for Cu(AMT-IDA) and Cu(CO-IDA) at various pHs. The measurements for each complex are averages from analyses of 2-6 experiments.

| Isothermal Titration Calorimetry Results |  |  |  |  |  |
| --- | --- | --- | --- | --- | --- |
| Copper Complex | pH | $\Delta H$ (kcal/mol) | $\Delta H$<br>Standard<br>Dev. | $K_d$ | $K_d$ Standard Dev. |
| 1 mM Cu(Amt-IDA) | 8 | -5.8 | 0.3 | $0.87 \times 10^{-4}$ | $3.7 \times 10^{-5}$ |
| | 7 | -6.5 | 1.1 | $2.4 \times 10^{-4}$ | $8.3 \times 10^{-5}$ |
| | 6 | -5.5 | 1.1 | $1.5 \times 10^{-4}$ | $3.5 \times 10^{-5}$ |
| 1 mM Cu(CO-IDA) | 8 | -8.4 | 0.4 | $1.8 \times 10^{-4}$ | $2.4 \times 10^{-5}$ |
| | 7 | -6.6 | 0.4 | $1.1 \times 10^{-4}$ | $1.2 \times 10^{-5}$ |
| | 6 | -4.8 | 0.5 | $2.8 \times 10^{-4}$ | $1.9 \times 10^{-5}$ |

Cu(CO-IDA) also had block kinetics similar to Cu^2+^(aq) and Cu(AMT-IDA), with nearly complete block at 0.50 and 1.0 mM by 25 minutes (Figure 2e) and no significant wash-out after 5 minutes (Figure 2f). Like Cu(AMT-IDA), unbinding from C2 (*k_3_*) is higher than for Cu^2+^(aq), as expected from ligand-induced destabilization. Also, entry (*k1*) is remarkably higher than for Cu(II) or Cu(AMT-IDA), suggesting that the flexible ligand enhances entry into the channel. But these variations are compensated by other changes, such that *K_d,eff_* is 101 nM, intermediate between the values for Cu(AMT-IDA) and Cu^2+^(aq). Again, the value suggests formation of a weak covalent bond between the CO-IDA complexed Cu(II) and the channel.

The outer dissociation constant (*K_d1_*) for Cu(AMT-IDA) bound to the initial site, i.e. dissociating from state C1, is 2.86 × 10^−4^ M^−1^, on the same order as the dissociation constant for AMT from M2 S31N, which can be calculated from the rate constants as 1.1 × 10^−4^ M^−1^ (19). The association rate constant, *k_1_*, is ~5x higher for AMT than for Cu(AMT-IDA) and the dissociation rate constant, *k_4_*, ~2 higher, suggesting that the Cu-diacetate adduct inhibits passage through the Val27 stricture at the channel entry compared to AMT without the adduct. Similarly, cyclooctylamine, which is known to inhibit influenza A M2 somewhat better than AMT (24), gives faster entry and exit rate constants for Cu(CO-IDA) than AMT with a net lower *K_d1_* (Table 2), as would be expected because of the flexibility of its alkyl ring. The failure of Cu(IDA), with no hydrophobic adduct, to block the channel (Figure S4) demonstrates that the hydrophobic moiety in the ligand is necessary. Therefore, we propose that the first binding step can be viewed as entry into the channel facilitated by the hydrophobic moiety associating with the hydrophobic Val27 cluster. Although binding is weak in the M2 S31N, it is still sufficient to present the Cu(II) to the covalent bond acceptor in the channel. It is then followed by covalent binding of the ligated Cu(II). The kinetics and affinity of the second site confirm that the covalent bond acceptor in the channel is the His37 cluster.

### Electrophysiology – S31N-G34E

The S31N-G34E variant has glutamates in the place of glycines in the channel, which could potentially create steric hindrance preventing the copper complexes from reaching the His37 cluster. Blocking and wash-out traces for M2 S31N-G34E exposed to 100 μM of Cu^2+^(aq), Cu(AMT-IDA), or Cu(CO-IDA) were obtained using TEVC (Figure 3). Cu^2+^(aq) (Figure 3a) shows blocking kinetics similar in the S31N-G34E M2 to those in the S31N M2 (Figure 2a,b), with an initial fast phase in the wash-in and modest recovery in the wash-out. Cu(CO-IDA) blocks 86% of the M2 current after 12 minutes of perfusion (Figure 3b), more than double the time to block M2 by Cu^2+^(aq). Cu(CO-IDA) shows more wash-out compared to Cu^2+^(aq) (Figure 3a), though the appearance is exaggerated due to timescale contraction. Similarly, Cu(AMT-IDA) blocks 87% of the M2 current after 10 minutes of perfusion (Figure 3c). Again, the wash-out is modest. The block kinetics of S31N-G34E by AMT contrast sharply to those of the metal complexes. Maximal block by 100 μM AMT is reached quickly, but the total block is only 30% (Figure 3d). This result is similar to previous results with AMT in M2 S31N (19). The wash-out is nearly complete after 3 minutes of perfusion with Barth’s pH 5.3 minus AMT. Thus, AMT is not effective in blocking this double AMT-resistance mutant, as would be expected. These results show that the metal complexes block M2 current even when glutamates replace the glycines in the channel.

**Figure 3:**
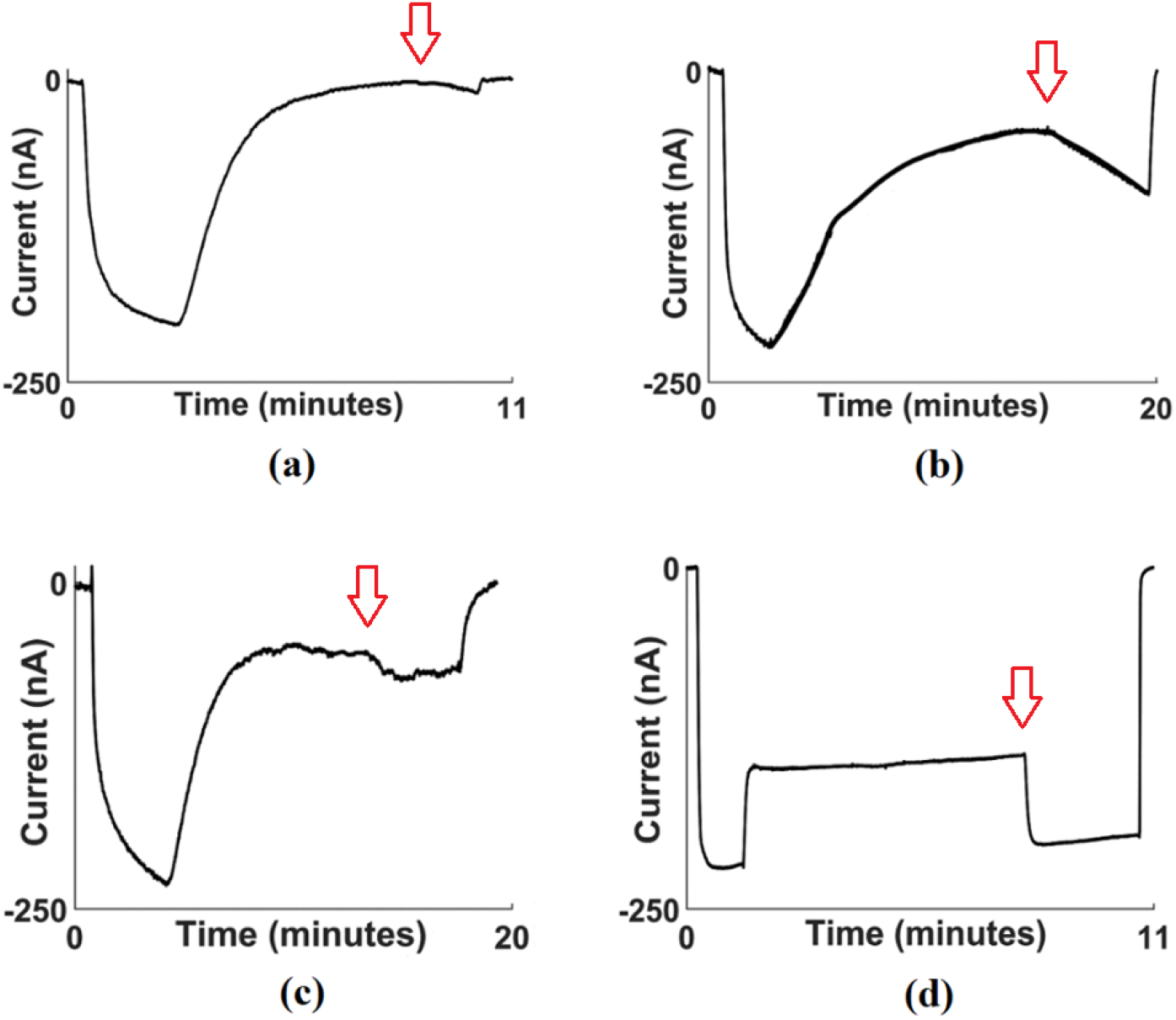
TEVC current traces for oocytes transfected with S31N-G34E M2. Representative traces show whole cell currents tested by perfusion in 0.1 mM solutions of a) Cu2+(aq) b) Cu(CO-IDA) c) Cu(AMT-IDA) d) AMT, which starts at 1-2.5 mins at the time of onset of block of inward current. Washout with Barth’s pH 5.3 without blocker begins at the red arrow. Perfusion is returned to the starting solution, Barth’s pH 7.4, at the end of the experiment

### Isothermal Titration Calorimetry

In order to evaluate the binding strength of the copper complexes with the imidazoles of the His37 cluster observed in electrophysiology, ITC measurements were performed (see representative temperature plots and area-under-the-curve data in Figures S2-S4). When imidazole was added to solutions of Cu(Amt-IDA), the enthalpy values at pH levels of 6, 7, and 8 were similar to each other, with values of −5.5 ± 1.1 kcal/mol (pH =6), −6.5 ± 1.1 kcal/mol (pH = 7), and −5.8 ± 0.3 kcal/mol (pH = 8) (Table 3). The enthalpy values of imidazole binding to Cu(CO-IDA) became more negative as pH increased, with values of −4.8 ± 0.5 kcal/mol (pH 6), −6.6 ± 0.4 kcal/mol (pH 7), and −8.4 ± 0.4 kcal/mol (pH 8). Both complexes are comparable at pH 6 and 7, the only different at pH of 8, with the Cu(CO-IDA) having the most negative enthalpy of binding. However, there is an interesting difference in binding illuminated when the pH varies. At pH 6 and 7, Cu(AMT-IDA) and Cu(CO-IDA) have approximately the same enthalpy of binding with imidazole (−5.5 vs −4.8 and −6.5 vs −6.6). Yet, at pH 8 there is a difference of 2.6 kcal/mol between the copper complexes binding imidazole. Both copper complexes have *<u>K</u>_<u>d</u>_* values of the same magnitude (10^−4^ M) suggesting strong binding for both copper complexes to imidazole. The enthalpy values and equilibrium constants for Cu(AMT-IDA) and Cu(CO-IDA) binding to imidazole are comparable to imidazole binding to aqueous Cu^2+^ with enthalpy = −7.2 kcal/mol and *<u>K</u>_<u>d</u>_* = 4.9 × 10^−5^ (25).

### Quantum Chemical Modeling of Imidazole Coordination to Copper Complexes

Table 4 reports the ωB97X-D3 calculated imidazole coordination energies to Cu(CO-IDA)(OH_2_)_2_ and Cu(AMT-IDA)(OH_2_)_2_ for the reactions in Figure 4. This model represents the enthalpy estimate (with inclusion of solvation free energy) of coordination at pH 7 where the imidazole is de-protonated. The calculated Δ*H* value (at 298 K at 1 atm (no concentration correction was applied)) for Cu(CO-IDA) is –12.45 kcal/mol (Table 4), which is reasonably close to the measured enthalpy of −6.6 kcal/mol (Table 3), especially considering only two explicit waters were used in the calculation. The calculated Cu-imidazole nitrogen bond length is 2.03 Å, which is in the range of 1.97-2.16 Å found in crystal structures (26–28).

**Figure 4:**
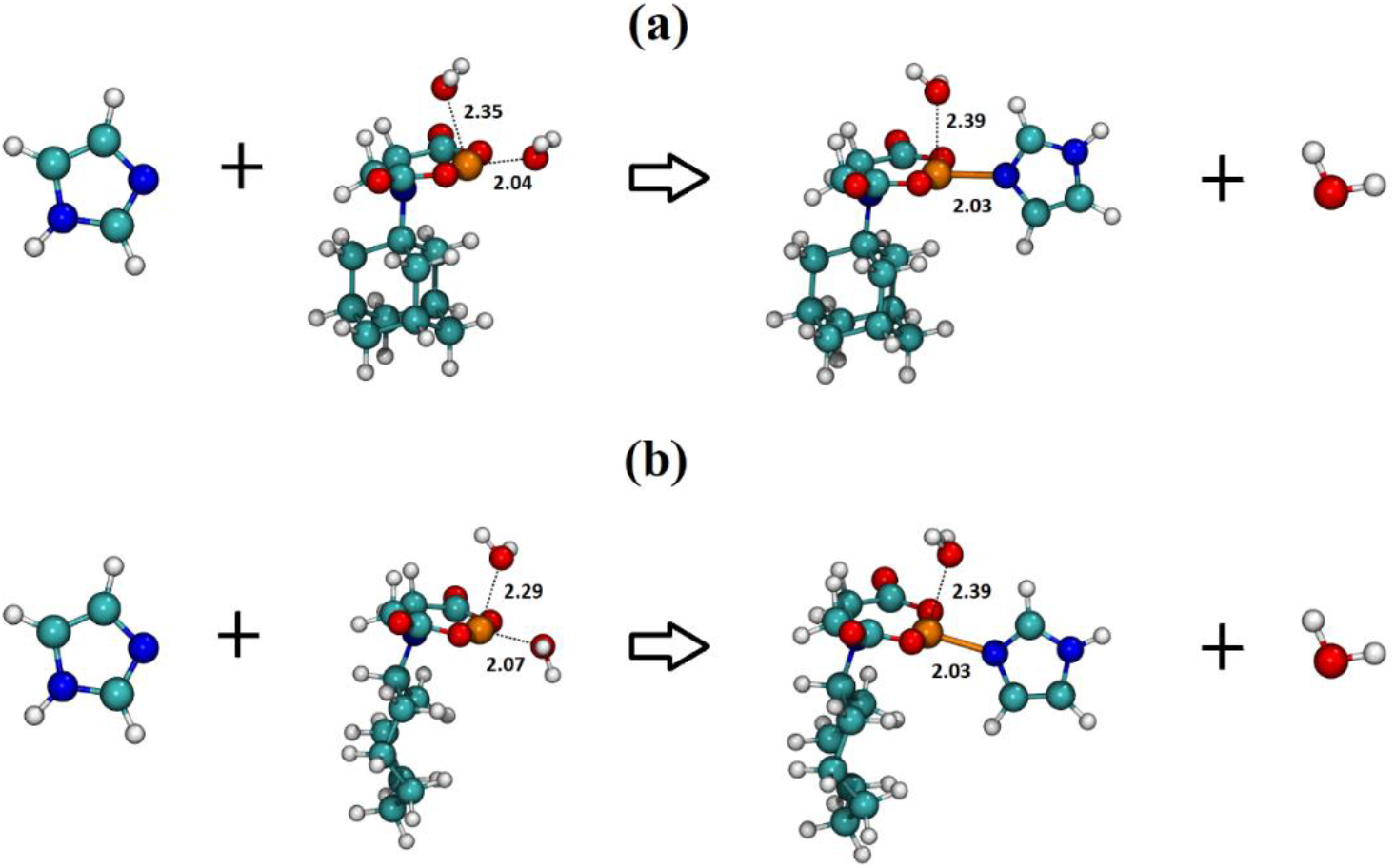
Cu(AMT-IDA) (a) and Cu(CO-IDA) (b) computational model reaction with unprotonated imidazole in implicit water solvent.

**Table 4:**
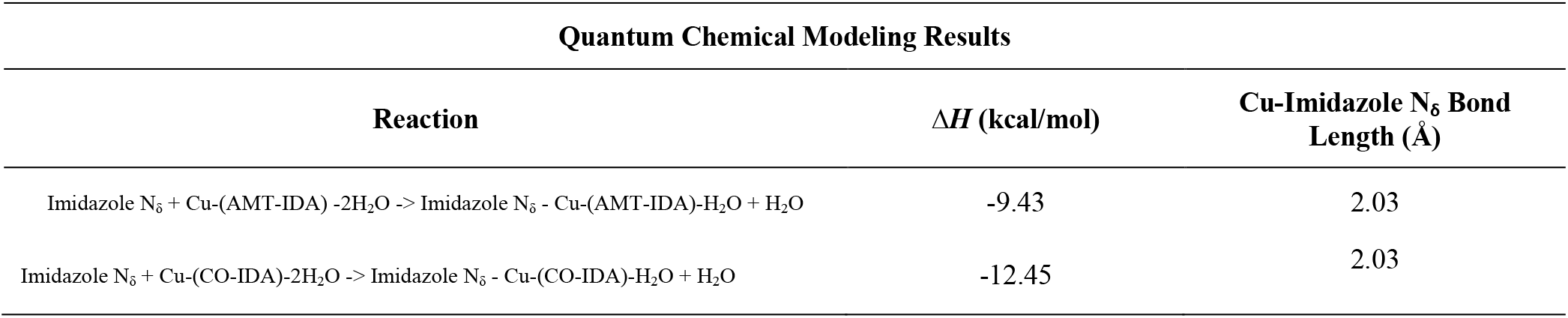
Copper complexes binding imidazole N_δ_ in implicit water solvent. Calculated enthalpy of reaction (difference between heat of formation of the reactants and products) using the ωB97X-D3 DFT functional in Terachem.

The calculated imidazole coordination enthalpy for Cu(AMT-IDA) is –9.43 kcal/mol (Table 4), and this value is reasonably close to the measured enthalpy value of −6.5 kcal/mol (Table 3). The calculated Cu-imidazole nitrogen bond length is 2.03 Å. These DFT coordination energies suggest that the ITC values are measuring imidazole substitution for water at Cu in the equatorial position and in the plane of the IDA ligand. Enthalpy calculations were done for a model with a second imidazole binding in the axial position on the copper for both complexes (Table S1 and Figure S6). The calculated enthalpy for two imidazole coordination with Cu(AMT-IDA) was − 15.79 kcal/mol and −18.37 kcal/mol for Cu(CO-IDA). Though the model suggests two imidazoles could bind to the copper complexes, the second imidazole binding is weaker than the first imidazole.

### Quantum Chemical Model – Copper Complexes Binding His37 In S31N M2 and its G34E homolog

The geometry optimization of Cu(CO-IDA) in the M2 S31N variant channel shows the nitrogen is bonded in an equatorial position on the copper while the copper remains complexed with the CO-IDA (Figures 5a and 5b). The bond length between the copper and imidazole nitrogen is 1.99 Å. The Cu-N_Imid_ bond directionality is in the copper’s remaining equatorial position, due to the equatorial position providing a stronger bond than an axial position. Indeed, it appears that tridentate complexation of Cu(II) is optimal for this situation (see *Summary of related projects* in Supplemental Information). The binding of two imidazoles to the copper (one equatorial the second axial) was attempted in these simulations. However, no proper starting geometries were achievable without extreme rotation of the copper complex and distortion of the M2 backbone. For more than one imidazole to bind the copper, it appears that the copper would need to dissociate from the ligand. The geometry optimization structure for Cu(AMT-IDA) in the M2 S31N variant is shown in Figures 5c, and 5d. The optimized structure result is similar to Cu(CO-IDA). The adamantyl group is not as flexible as the cyclooctyl, but it still fits down near the His37 cluster allowing the copper to bind to an imidazole nitrogen. The imidazole nitrogen binds in an open equatorial position of the copper while the copper remains complexed with the AMT-IDA. The Cu-N_Imid_ bond length is 2.07 Å. The geometry optimization structure for Cu(CO-IDA) in the M2 S31N-G34E variant shows the glutamates at position 34 do not prevent Cu(CO-IDA) from reaching a position to bind with a His37 imidazole (Figures 6a and 6b). This agrees with the electrophysiology results that Cu(CO-IDA) still blocks M2 S31N-G34E current. The geometry optimization of Cu(AMT-IDA) in the M2 S31N-G34E variant also showed that the complex fits near the His37 cluster and the copper bonds to an imidazole nitrogen (Figures 6c and 6d).

**Figure 5:**
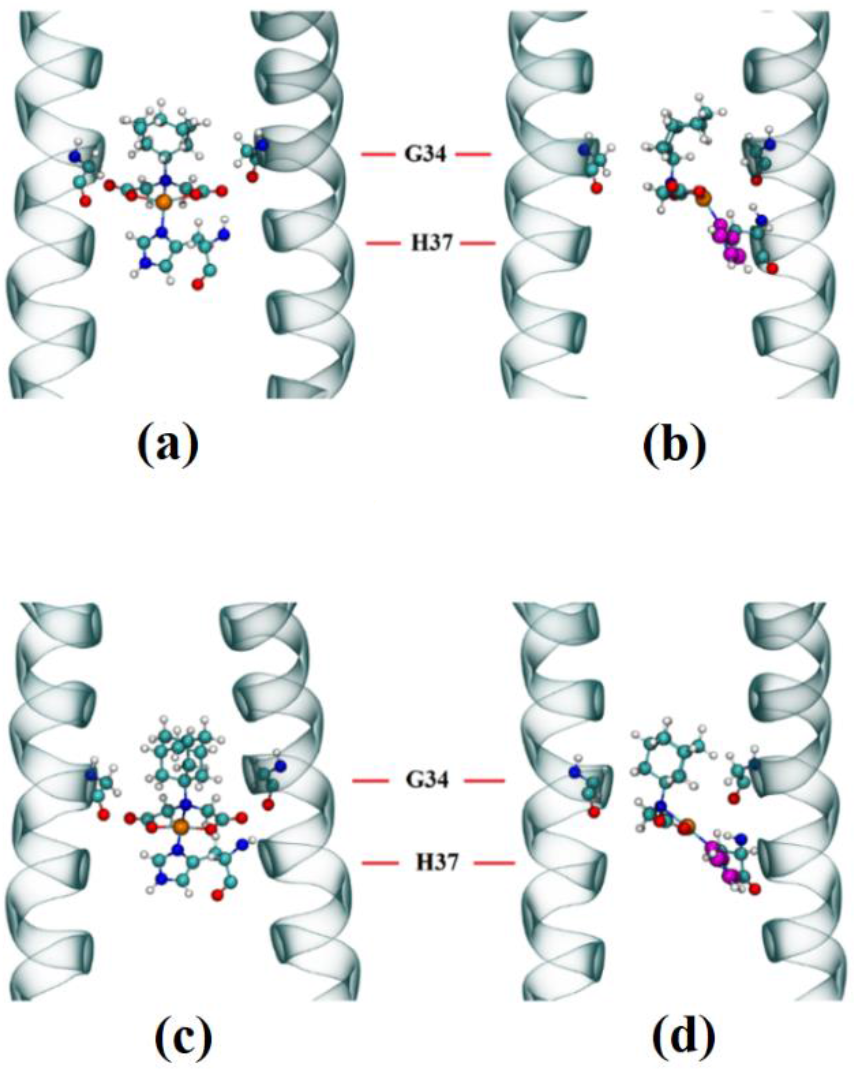
Computational chemistry geometry optimization of Cu(CO-IDA) (1) and Cu(AMT-IDA) (2) in M2 S31N. The simulation was done in gas phase. The ribbons represent the backbone of M2. Cyan atoms are carbons, white atoms are hydrogens, Red atoms are oxygens, blue atoms are nitrogens, and the orange atom is copper. a) Face-on view. The front and back ribbon is not displayed for easier visualization of the imidazole binding with the copper complex, but the histidine is still connected to the backbone as shown in the side view. b) Side view. Magenta used to highlight imidazole carbons and nitrogens.

**Figure 6:**
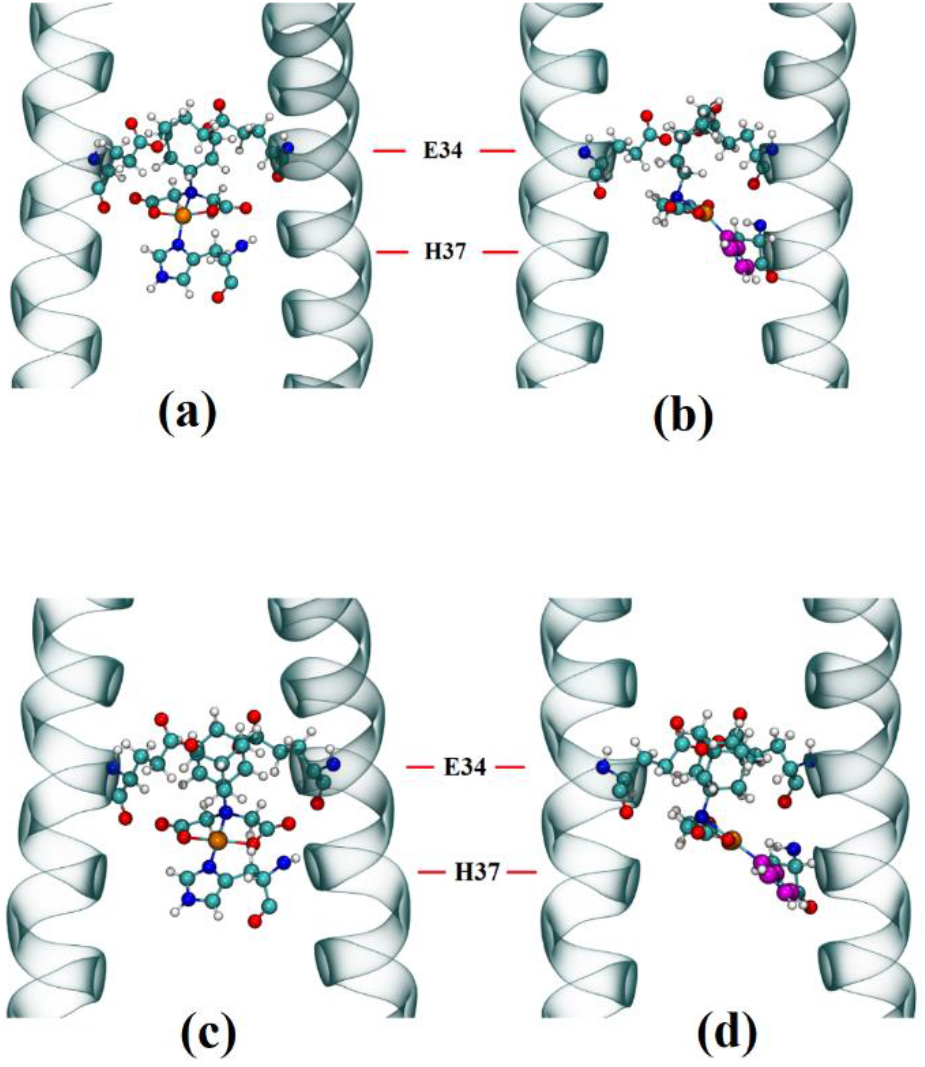
Computational chemistry geometry optimization of Cu(CO-IDA) (1) and Cu(AMT-IDA) (2) in M2 S31N-G34E. The simulation was done in the gas phase. The ribbons represent the backbone of M2. Cyan atoms are carbons, white atoms are hydrogens, Red atoms are oxygens, blue atoms are nitrogens, and the orange atom is copper. a) Face-on view. The front and back ribbon is not displayed for easier visualization of the imidazole binding with the copper complex, but the histidine is still connected to the backbone as shown in the side view. b) Side view. Magenta used to highlight imidazole carbons and nitrogens.

In summary, the ITC and computational chemistry results provide insight into the ability of the copper complexes to fit in the M2 channel near the His37 cluster and deliver the copper to the imidazole nitrogen. This provides an explanation for the two-stage M2 current block in the electrophysiology results, and the invulnerability of the complexes to resistance formation (17). Considering that IDA ligation of the copper substantially reduces its toxicity in zebrafish embryos (17), it would be interesting to examine toxicity and ADME properties of the complexes in higher animal models.

## Conclusion

Cu(CO-IDA) and Cu(AMT-IDA) bind to the M2 channel in S31N and S31N-G34E. A two-site mathematical model used in a global fit with the electrophysiology traces varying in copper complex concentration provided rate constants for Cu(CO-IDA) and Cu(AMT-IDA) binding in M2 S31N. The copper complexes get in to the His37 binding site at nearly the same rate as Cu^2+^(aq). The Cu^2+^(aq) and copper complexes are slow to leave the His37 binding site as shown by the dissociation rate constants being much smaller than the association rate constants, which implies strong binding to the His37 site. The copper complexes reside longer in the M2 channel than Cu^2+^(aq). The independent fit from ITC shows that the copper complexes are capable of binding one imidazole strongly, which is in agreement with the quantum chemical model. The quantum chemical model of the copper complexes in the channel show that the complexed copper binds to one imidazole in both the S31N and S31N-G34E M2 variants.

## Acknowledgments

The authors thank Jonathan Lynch for technical help in synthesizing the compounds, Thomas Walker for help performing ITC experiments, and Dr. Jason Kenealy for guidance on and use of his ITC instrument. This project was supported by funding from Brigham Young University to KM, DB, and RH. The Fulton Supercomputing Lab in the BYU Office of Research Computing provided computer resources. D.H.E acknowledges funding from the National Science Foundation, CHE-1952420.

## References

1. Hay, A. J., A. J. Wolstenholme, J. J. Skehel, and M. H. Smith. 1985. The molecular basis of the specific anti-influenza action of amantadine. The EMBO Journal 4(11):3021–3024.

2. Balannik, V., J. Wang, Y. Ohigashi, X. Jing, E. Magavern, R. A. Lamb, W. F. Degrado, and L. H. Pinto. 2009. Design and pharmacological characterization of inhibitors of amantadine-resistant mutants of the M2 ion channel of influenza A virus. Biochemistry 48:11872–11882.

3. Zhao, X., Y. Jie, M. R. Rosenberg, J. Wan, S. Zeng, W. Cui, Y. Xiao, Z. Li, Z. Tu, M. G. Casarotto, and W. Hu. 2012. Design and synthesis of pinanamine derivatives as anti-influenza A M2 ion channel inhibitors. Antivir Res 96(2):91–99.

4. Wang, J., C. Ma, H. Jo, B. Canturk, G. Fiorin, L. H. Pinto, R. A. Lamb, M. L. Klein, and W. F. Degrado. 2013. Discovery of novel dual inhibitors of wild-type and the most prevalent drug-resistant mutant, S31N, of M2 proton channel from influenza A virus. Journal of medicinal chemistry 56:2804–2812.

5. Rey-Carrizo, M., E. Torres, C. Ma, M. Barniol-Xicota, J. Wang, Y. Wu, L. Naesens, W. F. DeGrado, R. A. Lamb, L. H. Pinto, and S. Vazquez. 2013. 3-Azatetracyclo[5.2.1.1(5,8).0(1,5)]undecane derivatives: from wild-type inhibitors of the M2 ion channel of influenza A virus to derivatives with potent activity against the V27A mutant. Journal of medicinal chemistry 56(22):9265–9274.

6. Rey-Carrizo, M., M. Barniol-Xicota, C. Ma, M. Frigole-Vivas, E. Torres, L. Naesens, S. Llabres, J. Juarez-Jimenez, F. J. Luque, W. F. DeGrado, R. A. Lamb, L. H. Pinto, and S. Vazquez. 2014. Easily accessible polycyclic amines that inhibit the wild-type and amantadine-resistant mutants of the M2 channel of influenza A virus. Journal of medicinal chemistry 57(13):5738–5747.

7. Kolocouris, A., C. Tzitzoglaki, F. B. Johnson, R. Zell, A. K. Wright, T. A. Cross, I. Tietjen, D. Fedida, and D. D. Busath. 2014. Aminoadamantanes with persistent in vitro efficacy against H1N1 (2009) influenza A. Journal of medicinal chemistry 57(11):4629–4639.

8. Wu, X., X. Wu, Q. Sun, C. Zhang, S. Yang, L. Li, and Z. Jia. 2017. Progress of small molecular inhibitors in the development of anti-influenza virus agents. Theranostics 7(4):826–845.

9. Wang, Y., Y. Hu, S. Xu, Y. Zhang, R. Musharrafieh, R. K. Hau, C. Ma, and J. Wang. 2018. In Vitro Pharmacokinetic Optimizations of AM2-S31N Channel Blockers Led to the Discovery of Slow-Binding Inhibitors with Potent Antiviral Activity against Drug-Resistant Influenza A Viruses. Journal of medicinal chemistry 61(3):1074–1085.

10. Musharrafieh, R., P. Lagarias, C. Ma, R. Hau, A. Romano, G. Lambrinidis, A. Kolocouris, and J. Wang. 2020. Investigation of the Drug Resistance Mechanism of M2-S31N Channel Blockers through Biomolecular Simulations and Viral Passage Experiments. ACS Pharmacology & Translational Science https://dx.doi.org/10.1021/acsptsci.0c00018.

11. Musharrafieh, R., C. Ma, and J. Wang. 2020. Discovery of M2 channel blockers targeting the drug-resistant double mutants M2-S31N/L26I and M2-S31N/V27A from the influenza A viruses. European journal of pharmaceutical sciences: official journal of the European Federation for Pharmaceutical Sciences 141:105124.

12. Gordon, N. A., K. L. McGuire, S. K. Wallentine, G. A. Mohl, J. D. Lynch, R. G. Harrison, and D. D. Busath. 2017. Divalent copper complexes as influenza A M2 inhibitors. Antivir Res 147:100–106.

13. Chizhmakov, I. V., F. M. Geraghty, D. C. Ogden, A. Hayhurst, M. Antoniou, and A. J. Hay. 1996. Selective proton permeability and pH regulation of the influenza virus M2 channel expressed in mouse erythroleukaemia cells. Journal of Physiology 494 (Pt 2):329–336.

14. Rannulu, N. S., and M. T. Rodgers. 2005. Solvation of copper ions by imidazole: structures and sequential binding energies of Cu+(imidazole)x, x = 1-4. Competition between ion solvation and hydrogen bonding. Physical Chemistry Chemical Physics: PCCP 7(5):1014–1025.

15. Gandhi, C. S., K. Shuck, J. D. Lear, G. R. Dieckmann, W. F. DeGrado, R. A. Lamb, and L. H. Pinto. 1999. Cu(II) inhibition of the proton translocation machinery of the influenza A virus M2 protein. Journal of Biological Chemistry 274(9):5474–5482.

16. Su, Y., F. Hu, and M. Hong. 2012. Paramagnetic Cu(II) for probing membrane protein structure and function: inhibition mechanism of the influenza M2 proton channel. Journal of the American Chemical Society 134(20):8693–8702.

17. McGuire, K. L., J. Hogge, A. Hintze, N. Liddle, N. Nelson, J. Pollock, A. Brown, S. Facer, S. Walker, R. G. Harrison, and D. D. Busath. 2019. Copper Complexes as Influenza Antivirals: Reduced Zebrafish Toxicity. Noble Metals in Medicine DOI 10.5772/intechopen.88786:1-15.

18. Mash, H. E., Y. P. Chin, L. Sigg, R. Hari, and H. Xue. 2003. Complexation of copper by zwitterionic aminosulfonic (Good) buffers. Anal Chem 75(3):671–677.

19. McGuire, K. L., J. T. Hill, and D. D. Busath. Increased dissociation of adamantanamines in Influenza A M2 S31N with partial block by rimantadine. Submitted.

20. Bevington, P. R. 1969. Data Reduction and Error Analysis for the Physical Sciences. McGraw Hill, New York.

21. Grimme, S. 2006. Semiempirical GGA-type density functional constructed with a long-range dispersion correction. Journal of Computational Chemistry 27(15):1787–1799.

22. Klamt, A., and G. Schuurmann. 1993. COSMO - A new approach to dielectric screening in solvents with explicit expresssions for the screening energy and its gradient. J. Chem. Soc. - Perkin Transactions 2(5):799–805.

23. Dapprich, S., I. Komaromi, K. S. Byun, K. Morokuma, and M. J. Frisch. 1999. A new ONIOM implementation in Gaussian 98. The calculation of energies, gradients and vibrational frequencies and electric field derivatives. Journal of Molecular Structure (Theochem) 462:1–21.

24. Lin, T. I., H. Heider, and C. Schroeder. 1997. Different modes of inhibition by adamantane amine derivatives and natural polyamines of the functionally reconstituted influenza virus M2 proton channel protein. J Gen Virol 78 (Pt 4):767–774.

25. Skenskaya, E. V., and M. K. Karapet’vants. 1966. Russian Journal of Inorganic Chemistry 11:1468.

26. Lin, D. D., and D. J. Xu. 2005. Synthesis and crystal structure of tetra(imidazole) copper(II) terephthalate. Journal of Coordination Chemistry 58(7):605–609.

27. Zhang, H. 2018. Crystal structure of *catena*-poly[chlorido-(μ_2_-chlorido)-bis(imidazole-κN)copper(II)] C_6_H_8_Cl_2_CuN_4_. Z. Kristallogr. 233(2):223–224.

28. Wen, W., X. Jimin, and X. Dongpo. 2010. Two copper complexes with imidazole ligands: Syntheses, crystal structures and fluorescence. Russian Journal of Inorganic Chemistry 55(3):384–389.

